# The full-length transcriptome of *C. elegans* using direct RNA sequencing

**DOI:** 10.1101/598763

**Authors:** Nathan P. Roach, Norah Sadowski, Amelia F. Alessi, Winston Timp, James Taylor, John K. Kim

## Abstract

Current transcriptome annotations have largely relied on short read lengths intrinsic to most widely used high-throughput cDNA sequencing technologies. For example, in the annotation of the *Caenorhabditis elegans* transcriptome, more than half of the transcript isoforms lack full-length support and instead rely on inference from short reads that do not span the full length of the isoform. We applied nanopore-based direct RNA sequencing to characterize the developmental polyadenylated transcriptome of *C. elegans*. Taking advantage of long reads spanning the full length of mRNA transcripts, we provide support for 20,902 splice isoforms across 14,115 genes, without the need for computational reconstruction of gene models. Of the isoforms identified, 2,188 are novel splice isoforms not present in the Wormbase WS265 annotation. Furthermore, we identified 16,325 3’ untranslated region (3’UTR) isoforms, 2,304 of which are novel and do not fall within 10 bp of existing 3’UTR datasets and annotations. Combining 3’UTRs and splice isoforms we identified 25,944 full-length isoforms. We also determined that poly(A) tail lengths of transcripts vary across development, as do the strengths of previously reported correlations between poly(A) tail length and expression level, and poly(A) tail length and 3’UTR length. Finally, we have formatted this data as a publically accessible track hub, enabling researchers to explore this dataset easily in a genome browser.

## Introduction

The nematode *Caenorhabditis elegans* is an ideal experimental model organism due to its compact, well-annotated genome (The C. elegans Sequencing Consortium 1998; Wilson 1999; Hillier et al. 2005; Gerstein et al. 2010), invariant cell lineage (Sulston et al. 1983), and wide-array of molecular methods. Our current understanding of the *C. elegans* transcriptome has been determined with EST libraries, cDNA based libraries, and Illumina-based cDNA and RNA sequencing (Walhout et al. 2000; Reboul et al. 2001; Lamesch et al. 2004; Hillier et al. 2009; Gerstein et al. 2010; Spieth et al. 2014; Tourasse et al. 2017). Most coding sequences (CDSs) span more than 600 nucleotides (excluding introns), and the typical *C. elegans* gene contains 6.4 coding exons on average (Spieth et al. 2014).

3’ untranslated regions (3’UTRs) are critically important features of mRNA transcripts that contain binding sites for RNA-binding proteins and small noncoding RNAs (Cai et al. 2009; Szostak and Gebauer 2013). Regulation of 3’ UTR length can therefore have profound impacts on mRNA expression, stability, and localization (Mayr and Bartel 2009; Andreassi and Riccio 2009; Kuersten and Goodwin 2003). Large-scale sequencing of the *C. elegans* 3’UTRs revealed median lengths of 130-140 nucleotides (nt) with an average length of ~211 nt (Jan et al. 2011; Mangone et al. 2010). In addition, poly(A) tails in *C. elegans* have a median length of approximately 57 nt at the L4 stage and short poly(A) tail lengths are a feature of highly expressed genes (Lima et al. 2017).

The average transcript in *C. elegans* is significantly longer than the maximum possible read length of Illumina sequencing. Therefore, current approaches to annotate the full length structure of the average *C. elegans* transcript isoform rely on manual curation of gene models based on a variety of data types, while more generally computational approaches to assemble transcript structures from bulk, short-read sequencing data utilize computationally expensive and imperfect inference (Williams et al. 2011; Spieth et al. 2014; Pertea et al. 2015; Trapnell et al. 2012). Calculating poly(A) tail lengths requires a sequencing approach capable of resolving long homopolymers, and determining 3’UTR structures requires an experimental or computational means of determining which reads reflect the 3’ most base included in the transcript before cleavage and polyadenylation. The specialized protocols and analyses used to measure poly(A) tail length and identify 3’UTRs with short read sequencing approaches cannot directly link these measurements to their splice isoform of origin, and in the case of 3’UTR identification instead rely on assigning putative cleavage sites to the nearest overlapping or upstream gene (Subtelny et al. 2014; Chang et al. 2014; Mangone et al. 2010; Jan et al. 2011; Blazie et al. 2017; Diag et al. 2018).

Nanopore sequencing, in contrast, has no theoretical upper limit to read length and is capable of sequencing transcripts from end to end at a single molecule level (Garalde et al. 2018; Jenjaroenpun et al. 2018; Workman et al. 2018). Nanopore based sequencing methods have been used to annotate transcriptome structure in a variety of organisms ranging from the relatively simple *Saccharomyces cerevisiae*, to complex human cell lines and cancers (Byrne et al. 2017; Bayega et al. 2018; Garalde et al. 2018; Jenjaroenpun et al. 2018; Tang et al. 2018; Volden et al. 2018; Workman et al. 2018; Kadobianskyi et al. 2019; Sessegolo et al. 2019). In nanopore-based direct RNA sequencing (dRNAseq), RNA reads are captured by the 3’ end of their poly(A) tail, and sequenced 3’ to 5’ natively, directly measuring the RNA molecule. The full length of the poly(A) tail is sequenced, and using a trained hidden Markov model, the length of the poly(A) tail for each read can be estimated (Workman et al. 2018). The 3’ most base in the alignment should reflect the true cleavage and polyadenylation site for the full transcript represented by that read, provided that base-calling, trimming of poly(A) and adapter sequences, and alignment had sufficient precision. Despite these advantages, adoption of dRNAseq and other nanopore based sequencing methods is hindered due to the technology’s high error rates, and the relative lack of bioinformatics tools and analysis pipelines designed for long error-rich reads.

In this study, we have generated an atlas of post-embryonic *C. elegans* transcript structure using dRNAseq to sequence RNA extracted from across its developmental life cycle. We provide full length support for previously annotated splice isoforms, as well as novel splice isoforms. Furthermore, we identify and characterize 3’UTRs and compare these to known datasets. We also estimate poly(A) tail lengths for our reads and examine global properties of these lengths across development. Finally, we have made this data available both in raw formats and as a custom track hub.

## Results

### Collection and sequencing of developmentally staged *C. elegans*

To capture the diversity of transcript isoforms expressed across *C. elegans* development, we created dRNAseq libraries in technical duplicates from larval stages L1 to L4, as well as young and mature hermaphrodite adults (Figure 1A) (Corsi et al. 2015). Because wild-type *C. elegans* exists largely as hermaphrodites with spontaneous males (<0.5%) emerging in the population through chromosome nondisjunction, we also obtained a male enriched sample using a *him-8* mutant that disrupts X chromosome segregation (Hodgkin et al. 1979; Broverman and Meneely 1994; Phillips et al. 2005). We further enriched for the male subpopulation by filtering them through a 35um mesh that allows the males to be collected in the filtrate.

**Figure 1.**
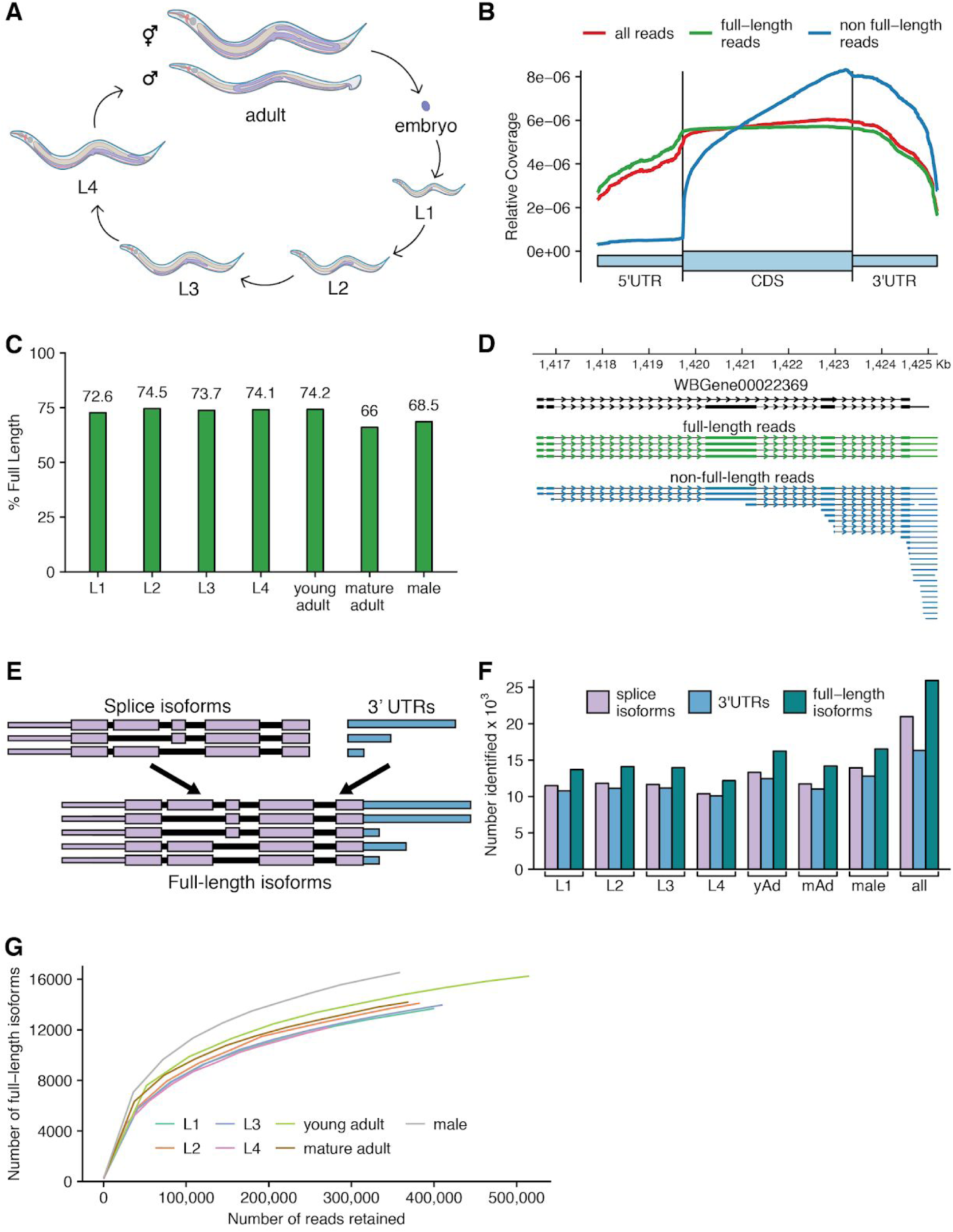
Overview of approach and sequencing of full-length isoforms. **(A)** Diagram of the C. elegans life cycle. **(B)** Plot of normalized coverage across the average coding gene with full-length (green), non-full-length (blue), and all reads (red) considered. **(C)** Percent of reads called full length in each stage. **(D)** Example locus showing reads aligning to the *WBGene00022369* locus (black). **(E)** Schematic defining “full-length isoform” as a combination of splice isoform and 3’UTR isoform. **(F)** Number of splice, 3’UTR, and full length isoforms observed across all stages. yAd = young adult, mAd = mature adult. **(G)** Saturation plot showing the number of full-length isoforms with support from one or more reads versus the number of reads considered, separated by stage. See Supplemental Figure 2 for equivalent plot with all stages combined.

Libraries were generated from RNA isolated by TriReagent (Ambion), poly(A) selected, and prepared for sequencing following the Oxford Nanopore Technologies SQK-RNA001 kit protocol with the exception of using Superscript IV (Thermo Fisher) in the optional reverse transcription step. The libraries were sequenced on an Oxford Nanopore Technologies GridION X5 (model #GRD-X5B002). Basecalling and adapter trimming of the reads was performed using poreplex (running albacore) (https://github.com/hyeshik/poreplex), resulting in over 540,000 reads that passed base calling quality control for each developmental stage sequenced, and 5.54 million total such reads (Supplemental Table 1). Reads had mean per base quality scores above 10 for each developmentally-staged sample, and median per base quality scores ranging between 9 and 10 for each sample. Median read lengths ranged between 573 and 687 for a given sample, while average read lengths were significantly longer, ranging from 739 to 934. Reads were aligned to the ce11 genome using minimap2, which successfully aligned 87.8% of our reads (Supplemental Table 2) (Li 2018).

### Identifying reads representing full length transcripts

While the majority of our reads correspond to full length transcripts (Figure 1B, C), a significant fraction of aligned reads failed to span the full length of an annotated transcript isoform; these reads were predominantly truncated relative to annotated isoforms at their 5’ ends, resulting in a 3’ bias in coverage from our total reads (e.g. Figure 1D). Including these reads in our downstream analysis would have artificially inflated the number of isoforms identified. Therefore, to make use of the long read lengths possible through dRNA-seq, reduce this 3’ bias, and eliminate the need to computationally reconstruct gene models, reads were split into ‘full-length’ and ‘non-full-length’ groups using existing coding sequence annotations, and only full-length reads were considered in downstream analyses (see Methods and Supplemental Figure 1 for an outline of the entire analysis). Note that nanopore sequencing reads are currently unable to capture the last 10-15 bases proximal to the 5’ end because of the structure of the pore-motor protein-RNA assembly, as reported previously (Workman et al. 2018)).

To determine the efficacy of this full-length filtering approach, we made an aggregate plot of normalized coverage across the average coding gene (Figure 1B). Supporting the validity of this approach, the non-full-length reads identified have a very extreme 3’ bias, while the full-length reads identified do not have the 3’ bias present in the total reads. Full length reads comprise the majority of reads in each dataset (Figure 1C). Combining all datasets, almost 2.9 million full length reads were obtained (Supplemental Table 2 for a breakdown of reads remaining after each filtering step).

In addition to full length filtering, a number of other filtering and analysis steps were applied, detailed in full in the methods. Briefly, reads were filtered if they 1) contained large insertions or large 3’ softclips (i.e. bases at the end of a read that fail to align); 2) had no detectable poly(A) tail signal; 3) had 5’ ends that weren’t aligned within 100 nt of an annotated transcript start site; 4) had a donor or acceptor splice site that couldn’t be assigned to an annotated donor or acceptor splice site (i.e., a splice site not within 15 bp of an annotated splice site); or 5) had retained introns. Following read filtering, the splice isoforms and 3’UTRs present in each developmental stage and across all stages were identified and reads were assigned to splice isoforms and 3’UTR isoforms as described in the methods.

### Identifying the full-length transcriptome

The full-length single-molecule resolution of nanopore sequencing means that, unlike short read sequencing, the full linear sequence of exons comprising a transcript and all of the associated splice junctions (i.e. the splice isoform) and the 3’UTR isoform are captured unambiguously together in a single read. This enables the identification of the “full-length transcriptome”, the set of full-length isoforms (splice isoform + 3’UTR isoform) observed together across all reads (Figure 1E). When considered across all developmental stages and conditions, 25,944 full-length isoforms were identified, comprised of 20,987 unique splice isoforms and 16,325 unique 3’UTRs (Figure 1F, Supplemental Table 3 for exact values). Over 12,000 full-length isoforms were identified in each stage. Because 3’UTRs were only called if there were 3 or more reads supporting the putative cleavage site, not all splice isoforms have an associated 3’UTR called, and therefore, some full-length isoforms have no high confidence 3’UTR call, and are in effect simply splice isoforms. This describes only a small number (3,518) of the full-length isoforms in the dataset.

To determine if these datasets were at or approaching saturation in the number of full-length isoforms identified, reads were randomly subsampled and the number of full-length isoforms that had support from one or more reads in the subsampled set was determined. These values were then plotted, and the relationship between the number of reads considered and the number of full-length isoforms supported was examined. As expected, none of the developmentally staged datasets appears to be saturated (Figure 1G). The number of isoforms identified across all stages appears to be approaching, but not at, saturation (Supplemental Figure 2). This implies that while further sequencing would expand the number of full-length isoforms identified in individual stages, it would likely have a modest effect on the total number of full-length isoforms identified across all stages.

The ability to resolve splice isoforms and 3’UTR isoforms together at single molecule resolution allows for identification of genes where the two features appear to be correlated. Notably, few examples of significant correlations between splice isoform use and 3’UTR isoform use were identified by Fisher exact test after multiple hypothesis testing correction (Supplemental Table 4). This is possibly due to lack of coverage, but more likely reflects an overall lack of coordination between splicing and polyadenylation site choice in *C. elegans*.

### Quantifying genes and splice-isoforms captured with full-length support

Less than half of the 30,133 isoforms with annotated introns in the WormBase WS265 annotation have full-length support (here full-length support means that every annotated intron in the isoform is supported by the same cDNA or EST) (Figure 2A) (Lee et al. 2018; WormBase web site 2018). By comparison, 17,658 full-length supported splice-isoforms across 13,622 genes were identified in our data, well above the 12,613 isoforms and 10,711 genes that have full-length support in the WormBase WS265 annotation. Comparing the genes and isoforms with full-length support in each dataset 4,234 genes and 7,404 isoforms were identified that did not previously have full-length support (Supplemental Figure 3A, Figure 2B). This dataset therefore significantly expands the number of *C. elegans* genes and isoforms supported by full-length reads.

**Figure 2.**
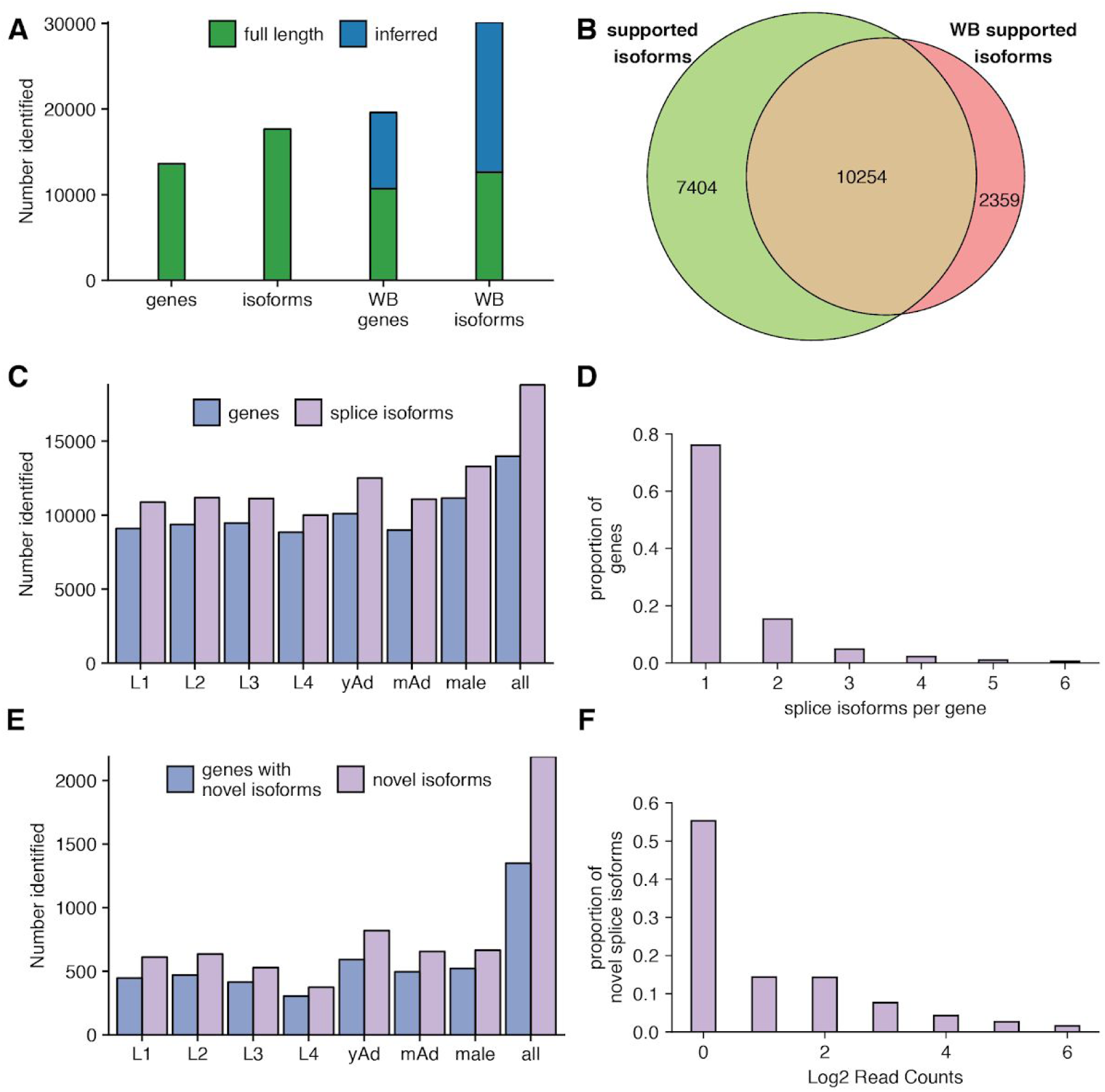
Capture of annotated and novel full-length splice isoforms. **(A)** Number of genes and isoforms captured with full length or inferred support in our dataset (left) versus the WormBase (WB) annotation (right). (WormBase web site 2018; Lee et al. 2018)**. (B)** Venn diagram of overlap between isoforms with full-length support in our dataset and those with full length support in the WormBase annotation. **(C)** Number of previously annotated splice isoforms and genes identified by our data across all stages. yAd = young adult, mAd = mature adult. **(D)** Density plot showing the number of splice isoforms identified per gene across our full dataset. **(E)** Number of novel isoforms and genes with novel isoforms identified across all stages. **(F)** Density plot showing the proportion of novel splice isoforms with a given number of reads supporting their structure.

To examine the changes of splice isoform usage in each developmental stage and across all stages, we plotted the number of previously annotated splice isoforms and genes observed in each stage (Figure 2C, Supplemental Table 3). We found more than 9,900 previously annotated splice isoforms in each stage, with males having the most identified genes and splice isoforms of any individual stage despite having fewer reads after our filtering steps than most other stages (Supplemental Table 2). Combining across all stages, over 18,000 splice-isoforms were observed. Most genes in our transcriptome data have only a single identified splice isoform, and the frequency of genes with a given number of isoforms decreases as the number of isoforms increases (Figure 2D), consistent with the WS265 annotation of the *C elegans* transcriptome (Supplemental Figure 3B) (Lee et al. 2018; WormBase web site 2018).

In addition to capturing previously annotated splice isoforms, the appeal of long-read single molecule sequencing is the ability to detect novel splice isoforms. To test our ability to identify novel splice isoforms after stringent filtering and splice site correction steps, we searched for isoforms with a set of splice junctions not present in the WormBase WS265 annotation. 2,188 novel splice isoforms were identified across all stages corresponding to 1,349 genes (Figure 2E) (Supplemental Table 3). Of these novel splice isoforms, 1,283 have novel splice junctions between annotated donor and acceptor splice sites and 173 have novel exons. To determine the level of support for these novel isoforms we generated a density plot showing the proportion of novel isoforms with a given number of reads supporting them (Figure 2F). The majority of identified novel splice isoforms were identified with only a single read supporting their structure, however almost 25% of these novel isoforms had 4 or more reads supporting them indicating that these are high confidence novel isoforms.

### Characterizing the identified 3’UTRome

Previous analyses of nanopore sequencing have largely centered on splice isoform identification and characterization while largely ignoring the 3’UTR. Because dRNAseq relies on sequencing in the 3’ to 5’ direction of mRNAs isolated by their poly(A) tails, full-length sequences of 3’UTRs are preferentially captured. After adapter trimming, discarding reads with large 3’ softclips, and realigning the 3’ softclipped portions of the remaining reads, we identified putative poly(A) cleavage sites and predicted stop codons to define full-length 3’UTRs. Using this method, 16,325 unique 3’UTR isoforms were identified, with over 10,000 3’UTRs identified in each stage (Figure 3A, Supplemental Table 3).

**Figure 3.**
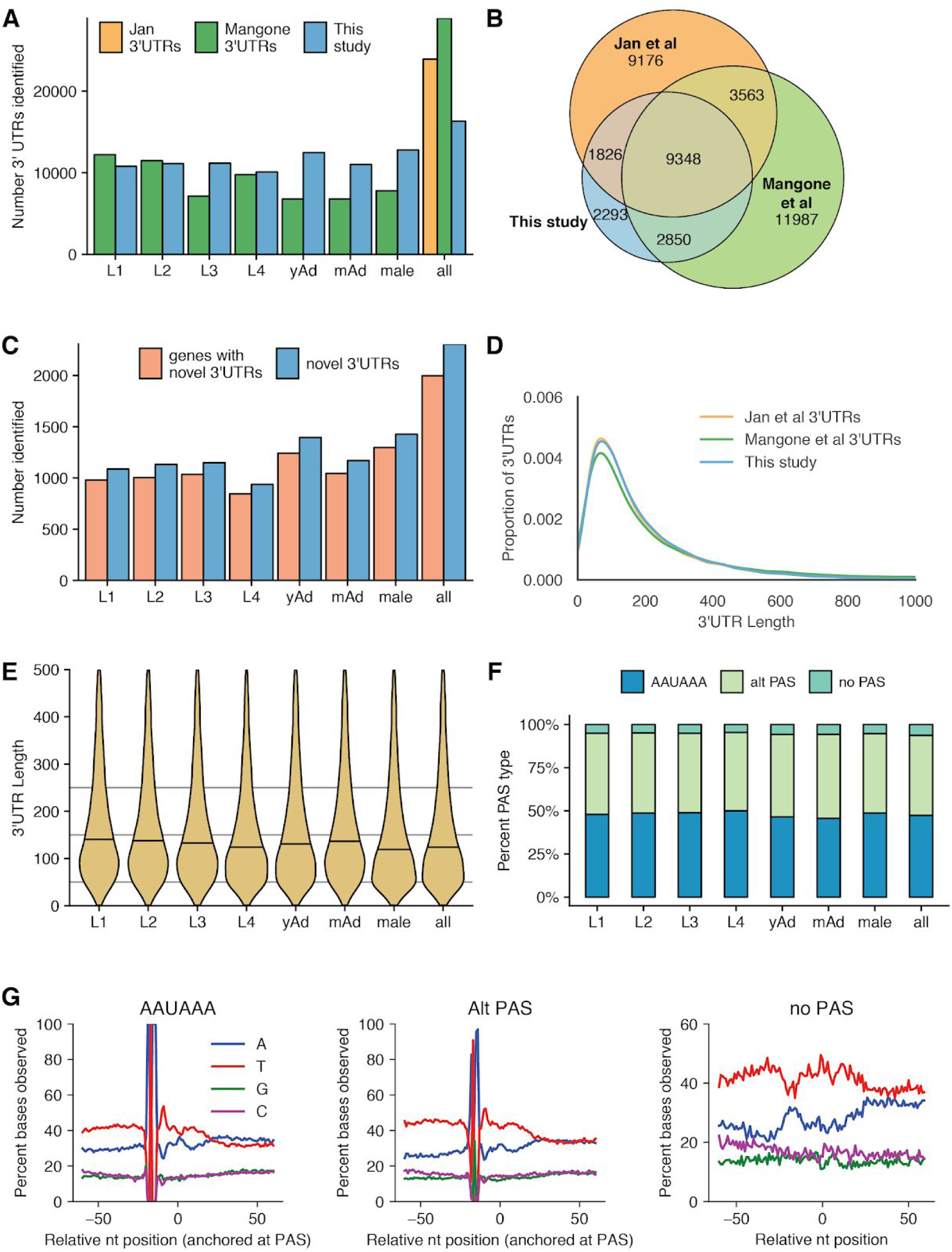
Properties of 3’UTRome. (A) Number of 3’UTRs observed across all stages, as compared to Mangone et al (Mangone et al. 2010) and Jan et al (Jan et al. 2011). yAd = young adult, mAd = mature adult. (B) Venn diagram showing overlap between 3’UTRs identified in this study, Jan et al, and Mangone et al. (C) Number of novel 3’UTRs and genes with novel 3’UTRs identified in each stage and across all stages. (D) Kernel density estimate plot of 3’UTR lengths from this study, Jan et al, and Mangone et al. (E) Violin plots showing 3’UTR length distributions across all stages. Horizontal black lines show the median of each stage.(F) Stacked bar chart showing percentage of UTRs with the specified polyadenylation signal (PAS) across all stages. (G) Nucleotide distributions around putative PAS sites and putative cleavage sites. Canonical PAS (AAUAAA) and alternative PAS (Alt PAS) distributions are anchored with the putative PAS hexamer at −19 nucleotides. The distribution of UTRs with no PAS is anchored with the putative cleavage site at 0.

To determine the accuracy of this 3’UTR calling method, we compared the 3’UTRs identified by this method with those from previously published datasets (including 3P-Seq and 3’RACE data) generated in *C. elegans* (Mangone et al. 2010; Jan et al. 2011). Of our identified 3’UTRs, 82.9% overlap with one or more of these 3’UTR datasets (Figure 3B). In addition, we identified 2,304 novel 3’UTRs that do not fall within 10 bp of existing 3’UTRs or WormBase 3’UTR annotations (Figure 3C). The 3’UTR length distribution in our data was nearly identical to those observed by Jan et al and Mangone et al (Figure 3D). In agreement with Mangone et al, our 3’UTR length distributions change over developmental stages, progressively decreasing from L1 through L4, and shorter in males than in hermaphroditic adults (Figure 3E). Curiously, the 3’UTR length distributions in adult stages were slightly longer than the length distribution of L4 3’UTRs in our datasets, in contrast to Mangone et al, which showed that adult 3’UTRs had a slightly shorter average 3’UTR length than L4.

Given 3’UTRs were shown to change length over development, we investigated whether PAS usage changed during development. Frequency of canonical and alternative PAS usage was quite consistent between adjacent developmental stages, although by chi squared tests there were significant differences in overall PAS usage between the L4 and young adult stage, as well as between hermaphroditic young adults and males (Figure 3F). Given that distribution of canonical and alternative PAS usage are consistent across the larval stages, where a significant shift in 3’UTR length distributions occurs, this suggests that 3’UTR length changes over development are largely independent of PAS usage.

As a final metric for the accuracy of this 3’UTRome, we plotted nucleotide distributions in windows around identified PAS sites and around putative cleavage sites (Figure 3G). This largely agrees with previously published nucleotide distributions in windows around identified PAS sites (Mangone et al. 2010). These distributions are AT-rich, with a peak in T frequencies just 3’ of the PAS site. It is possible that 3’UTRs identified by our method were inaccurate and broadly distributed around true cleavage sites, and by anchoring nucleotide distributions with putative PAS sites at −19 nucleotides the impact of these errors was eliminated. To test this possibility, we generated a density plot of the offsets of identified PAS sites from putative cleavage sites identified by our method, and found that these offsets were enriched close to the canonical −19 nucleotides from putative cleavage sites, indicating cleavage site calls from this method are accurate within a few base pairs (Supplemental Figure 4A & B).

Notably, at 3’UTR sites without a putative PAS identified, the nucleotide distribution observed lacks the enrichment of As in a window around the cleavage site noted in Mangone et al. Our method may be capturing a different set of 3’ UTRs with no PAS than the Mangone dataset did. Supporting this possibility, only 27% of the no PAS 3’UTRs in our dataset overlap with a Mangone et al 3’UTR, as compared with 73% of canonical PAS 3’UTRs in our data, and 65% of alternative PAS 3’UTRs in our data (Supplemental Figure 4C). In addition, the no PAS 3’UTRs that do overlap with a Mangone 3’UTR have a different nucleotide distribution than the no PAS Mangone 3’UTRs in general (Supplemental Figure 4D) (Mangone et al. 2010).

### Properties of poly(A) tail lengths

Poly(A) tails are known regulators of translation and transcript stability. However, profiling of poly(A) tail lengths at the transcriptome-wide level using short read sequencing is a relatively recent advance in the field (Subtelny et al. 2014; Chang et al. 2014; Lim et al. 2016). We have previously shown that, using a trained hidden Markov model, one can estimate the poly(A) tail length of dRNAseq reads using nanopolish (Workman et al. 2018). We performed these estimations on our datasets, providing a developmentally resolved poly(A) profiling dataset.

Global poly(A) tail length distributions are dynamic in the developing Drosophila melanogaster oocyte and embryo (Lim et al. 2016). To determine if there were comparable shifts in our poly(A) tail length distributions, we examined poly(A) tail lengths across the developmental stages in *C. elegans*. The poly(A) tail length distributions display only modest fluctuations, ranging from median values of 49 nt (L1) to 54 nt (L2) during larval development, although these shifts were considered to be statistically significant by Kolmogorov-Smirnov and Mann-Whitney U tests (Figure 4A). However, length distribution in all adult stages (young and mature hermaphrodites and males) are consistently longer than in the larval stages, with a median length of 58 in adults compared to an aggregate median length of 52 across all larval stages (p < 2.2e-16 by Kolmogorov-Smirnov and Mann-Whitney U tests). These data suggest that the most significant regulation of poly(A) tail lengths occurs between larval and adult stages during development.

**Figure 4.**
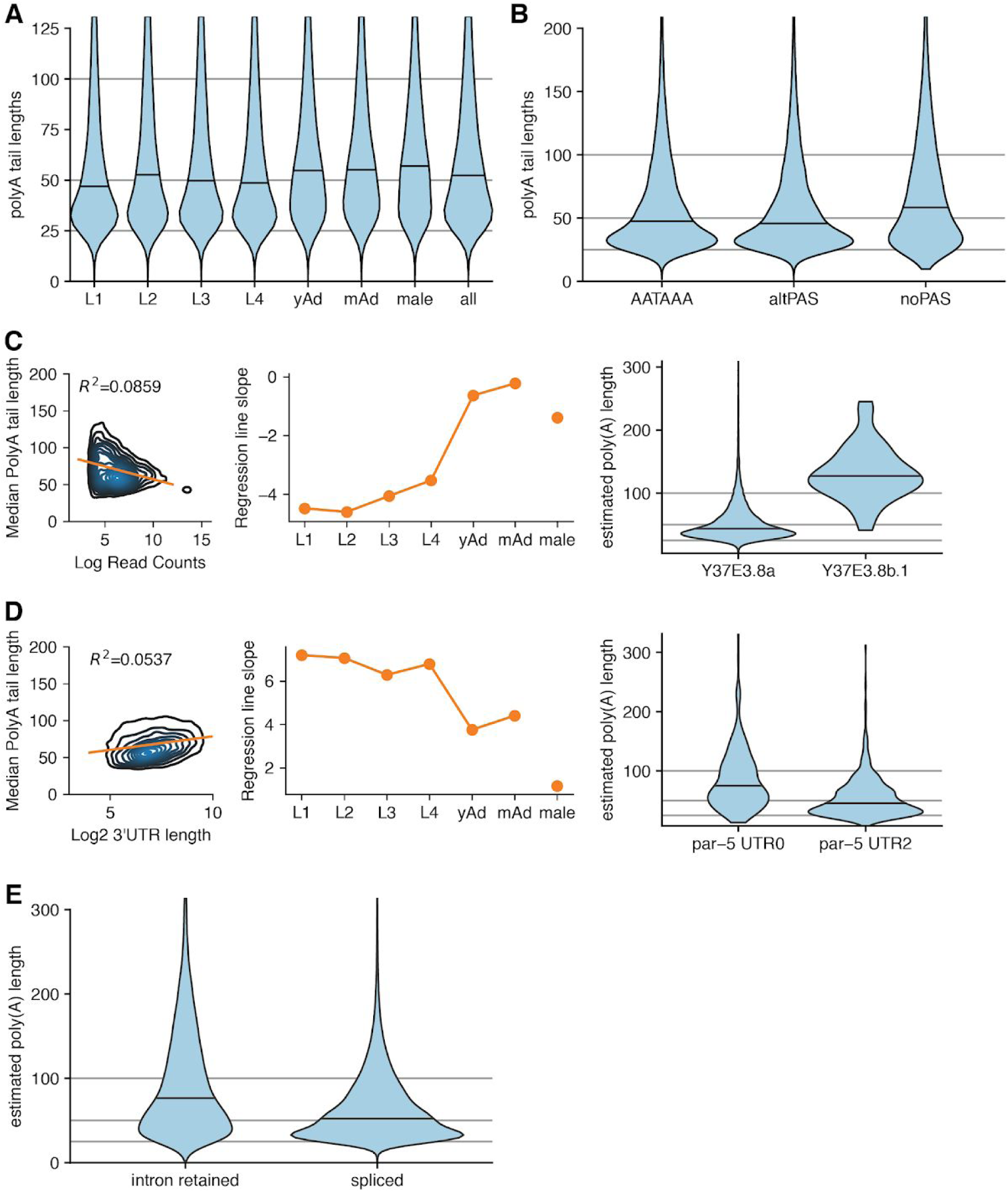
Properties of poly(A) tail length. **(A)** Violin plot of poly(A) tail length distributions across development. Horizontal black lines show the median of each stage. yAd = young adult, mAd = mature adult. **(B)** Poly(A) tail length distributions separated by the PAS type of the associated reads. **(C)** (left) Density plot showing correlation between poly(A) tail length and expression level by plotting median poly(A) tail length for each isoform versus the log of the expression level of that isoform (across all stages). Linear regression plotted in orange. (middle) Slope of linear regressions performed on median poly(A) tail length versus expression level data across developmental stage. (right) Example locus illustrating relationship between poly(A) tail length and expression level *Y37E3.8b.1* is lower expressed than *Y37E3.8a* with a longer poly(A) tail length distribution. **(D)** (left, middle) As in the left & middle panels of C, but instead plotting median poly(A) tail length versus the log of the 3’UTR length. (right) Example locus illustrating relationship between 3’UTR length and poly(A) tail length; *par-5* UTR0 is longer than *par-5* UTR2 and has a longer poly(A) tail length distribution. **(E)** Violin plots showing poly(A) tail length distributions in fully spliced versus intron-retention transcripts.

As a means of confirming the validity of our poly(A) tail length profiling approach, we compared our poly(A) estimates from the L4 stage with previously published poly(A) measurements from the L4 stage of *C. elegans* from mTAILseq (Lima et al. 2017). The length scale distributions of our L4 data and this dataset are quite similar, as both have peaks around 30-40 nt and extended spread toward the longer tail length range. (Supplemental Figure 5A). However, we did not identify the shoulder peaks present in the Lima et al dataset (Lima et al. 2017).

An advantage of profiling poly(A) tail lengths with dRNAseq versus short read sequencing is that poly(A) tail lengths are directly coupled to information about the splice isoforms and 3’UTR isoforms of the associated read. This allows comparisons and correlations between poly(A) tail lengths and aspects of transcript structure. One possible driver of differences in poly(A) tail lengths between reads could be that poly(A) tail length distributions may vary depending on whether the associated 3’UTR has a canonical PAS site. To test this possibility, we plotted poly(A) tail length distributions versus PAS type (i.e. canonical AAUAAA, alternative PAS, and no PAS) for reads from the L1 stage (Figure 4B). We find that all PAS types are significantly different from one another by Kolmogorov-Smirnov and Mann-Whitney-U tests (p < 2.2e-16), and 3’UTRs with no PAS have longer poly(A) tail lengths, on average, than poly(A) tails associated with either canonical and alternative PAS, with a median poly(A) tail length of 58 nt for 3’UTRs with no PAS, 46 nt for 3’UTRs with alternative PAS, and 48 nt for 3’UTRs with canonical PAS.

It has been reported that median poly(A) tail length and expression level are anticorrelated, such that highly expressed genes generally have shorter median poly(A) tail lengths (Lima et al. 2017; Legnini et al. 2019). To determine if this relationship holds in our datasets, we plotted the log of the number of reads supporting a given isoform versus the median poly(A) tail length for that isoform for transcripts with 10 or more reads supporting them (Figure 4C, left panel; Supplemental Figure 5B). A similar inverse correlation between median poly(A) tail length and number of reads supporting that isoform was observed in the L1 to L4 stages and when all stages were pooled (Supplemental Figure 5B). For example, the *a* isoform of the *Y37E3.8* gene is expressed much more than the *b.1* isoform (18,161 reads versus 38 reads), and has a significantly shorter poly(A) tail length distribution than the *b.1* isoform (Figure 4C, right panel). However, this correlation explains only a small fraction of the overall variation in the data, with the maximum R^2^ value of 0.1297. Interestingly, in the adult stages (both males and hermaphrodites), the slope of the regression lines between median poly(A) tail length and expression level were much more shallow, and the corresponding R^2^ values were much weaker with R^2^ values ranging from 0.0103 to 0.0004 (Figure 4C, middle panel; Supplemental Figure 5B). These results suggest that the inverse relationship between poly(A) length and expression level may vary depending on the developmental stage.

A recent study using FLAMseq, a PacBio sequencing method that also captures poly(A) tails and full length transcripts, demonstrated that poly(A) tail length and 3’UTR length were positively correlated (Legnini et al. 2019). Examining poly(A) tail length and 3’UTR lengths across all reads in our data, we also identify this same relationship (Figure 4D left panel). For example, the longer *par-5* 3’UTR isoform (termed 3’UTR 0; 486 nt) also has a longer poly(A) tail (median length 71) versus the shorter *par-5* 3’ UTR isoform (3’UTR 2; 51 nt) with a shorter poly(A) tail length distribution (median length 46) (Figure 4D right panel). However, the overall strength of this relationship also varies between developmental stages, and the slopes of the regression lines (and the corresponding R^2^ values) are smaller in adult stages than in larval stages (Figure 4D middle panel; Supplemental Figure 5C).

Finally, we examined the poly(A) tail length distributions between transcripts that are fully spliced versus those with retained introns. We previously showed in the human cell line GM12878 that intron retention correlates with transcripts with longer poly(A) tails (Workman et al. 2018). In our *C. elegans* datasets, we also found a positive correlation between intron retention and poly(A) tail length distributions by Kolmogorov-Smirnov and Mann-Whitney U tests, suggesting a conserved mechanism whereby nuclear transcripts possess longer poly(A) tails and supporting a model (Lima et al. 2017) in which poly(A) tails may be subject to post-transcriptional processing by deadenylation once exported into the cytoplasm.

### A public resource for full-length isoform information

To make our transcriptome dataset accessible to the research community, we have created a public custom track hub (https://bx.bio.jhu.edu/track-hubs/dRNAseq/hub.txt). This track hub contains the full-length filtered and non-filtered reads from each developmental stage, as well as the full-length isoforms supported across all stages at each locus. To ease access to this track hub, we registered it with the Track Hub Registry (https://www.trackhubregistry.org/). Users can therefore easily load this track hub in Ensembl (Zerbino et al. 2018) based genome browsers by searching public track hubs for “ce11 staged dRNAseq”. As a proof of the utility of this track hub, we loaded the track hub in the Ensembl genome browser and searched for *lin-14*, a gene with a well-studied 3’UTR that is subject to regulation by the *lin-4* microRNA (Wightman et al. 1991, 1993; Lee et al. 1993) but not currently annotated in the WormBase WS265 annotation (Lee et al. 2018; WormBase web site 2018). In our dataset, we identified the lin-14 3’UTR, as well as its splice isoforms, including a novel splice isoform (Figure 5A, “observed isoforms” track). As another example of the utility of this track hub, we searched for the locus *mlp-1*, a gene with multiple splice and 3’UTR isoforms identified, including multiple novel splice isoforms (isoforms 3, 5, and 7 of the observed isoform track in Figure 5B). These examples highlight possible uses of this resource by the research community to query currently unannotated 3’UTRs and splice isoforms.

**Figure 5.**
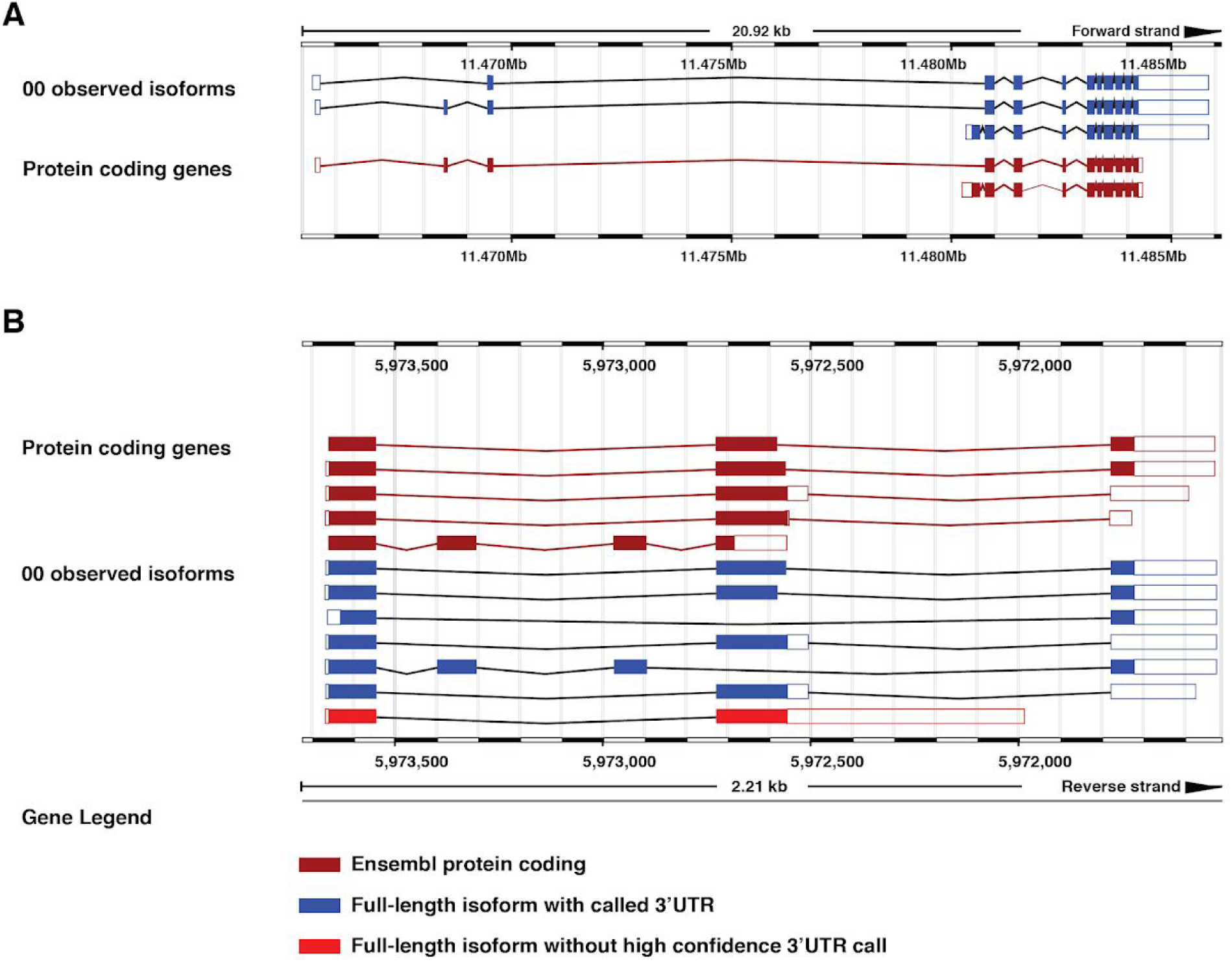
Examples highlighting utility of custom track hub. The *lin-14* **(A)**, or *mlp-1* **(B)** locus in the Ensembl genome browser including our custom track hub. Blue isoforms are full-length isoforms with an associated 3’UTR called, red isoforms have no high confidence 3’UTR called. Burgundy isoforms are protein coding models imported from WormBase.

## Discussion

Despite years of study, our understanding of the *C. elegans* transcriptome remains incomplete. Although studies have been performed profiling transcription start sites, splicing in both *cis* and *trans*, 3’UTR isoforms, poly(A) tail lengths, RNA base modifications, and gene and isoform expression levels, the short read lengths intrinsic to the prevailing technologies have been limited to examining one or two of these features at a time (Saito et al. 2013; Tourasse et al. 2017; Jan et al. 2011; Mangone et al. 2010; Lima et al. 2017; Zhao et al. 2015; Packer et al. 2019; Hillier et al. 2009). Even within these datasets, short read lengths and reliance on PCR amplification eliminate single molecule resolution, and make correlation of distant features within transcripts impossible. Although our study focuses primarily on splice isoforms, 3’UTR isoforms, and poly(A) tail lengths due to current limitations of nanopore sequencing technologies, in principle, modified approaches to dRNAseq would be capable of capturing all of the above features at a single molecule level.

Nanopore sequencing therefore poses both a unique set of opportunities and challenges that must be addressed in any analysis pipeline. The dRNAseq pipeline FLAIR (Full-length alternative isoform analysis of RNA) utilizes a hybrid sequencing approach in which matched short read sequencing is used to correct splice junctions in reads, and reads are clustered together in to splice isoforms if they share a common set of splice junctions (Tang et al. 2018).

We utilized an approach similar to that used by FLAIR, in which reads are corrected, in our case by an existing annotation, and clustered together by splice isoform. Our approach differs from FLAIR in several ways, including a full-length filtering step that reduces the impact of 3’ bias in our reads. A recent publication examining the utility of dRNAseq and cDNA nanopore sequencing to generate transcriptome annotations independently revealed that many nanopore sequencing reads fail to span the full-length of annotated transcript isoforms, highlighting the need for analysis pipelines that take the possibility of 5’ truncations into account in isoform identification (Soneson et al. 2019). Our full-length filtering approach partially addresses this concern, although, as noted by Soneson and colleagues, doing so reduces the number of usable reads, and likely impacts the quantitative nature of our data. A possible experimental approach to solving this problem could involve ligating a set of known nucleotides to the 5’ end of RNA transcripts after a decapping reaction, allowing for selection of full-length transcripts by filtering for reads flanked by signals corresponding to a poly(A) tail and the 5’ ligated product. This approach would incidentally also address the known problem that 10 - 15 nucleotides at the 5’ end of each strand are unable to be read (Workman et al. 2018).

Also distinguishing our approach from FLAIR is a novel means of calling 3’UTRs that has great utility in the generation of transcriptome annotations. Notably, we identify 3’UTR structures with a standard dRNAseq library preparation protocol meaning that, in principle, any dRNAseq experiment can be used to identify 3’UTRs using our method. The implications of this are potentially wide reaching, as experiments once used for comparative analysis of splice isoforms between conditions may now also be used in comparative analysis of 3’UTR isoforms.

By combining our 3’UTR and splice isoform calls, we identified almost 26,000 full-length transcript isoforms. It is likely that increased depth and additional sequencing of other developmental stages such as embryos and the stress-induced dauer stage would further increase the number of genes and isoforms identified, bringing this dataset closer to capturing the theoretical complete *C. elegans* transcriptome.

The ability to estimate poly(A) tail lengths for each read is another advantage of dRNAseq. Supporting the validity of our poly(A) profiling approach, the length distribution of the poly(A) tail length estimates we obtain in the L4 stage are quite similar to the distribution in the L4 stage reported by Lima et al, a study utilizing mTAILseq (Lima et al. 2017). Coupling of poly(A) tail lengths to aspects of 3’UTR structure and splice isoform allowed us to identify relationships between putative PAS sites and intron retention transcripts to poly(A) tail lengths. The relationship between PAS sites and poly(A) tail lengths is an interesting result that indicates there may be differential deposition or regulation of poly(A) tail length based on the presence or absence of an upstream PAS sequence. Longer poly(A) tails in intron retention transcripts could be indicative of intron retention transcripts being partially processed RNA still retained in the nucleus, as nuclear RNAs would be shielded from cytoplasmic deadenylase complexes. Neither of these relationships could be discovered by short read sequencing of poly(A) tails, demonstrating the efficacy of full-length single molecule sequencing.

A notable discovery of developmentally resolved poly(A) tail length profiling was the difference in features of poly(A) tail lengths between larval and adult stages. Overall poly(A) tail length distributions were longer in adult stages than in larval stages, and the strength of previously reported correlations between poly(A) tail lengths and expression level and poly(A) tail lengths and 3’UTR lengths were weaker in adult stages than larval stages. One possible explanation for these differences is the development of a functional germline in adult stages. In hermaphrodites, the cytoplasmic polyadenylases *gld-2* and *gld-4* are known to be active in the germline (Nousch et al. 2017; Suh et al. 2006; Schmid et al. 2009; Millonigg et al. 2014). Given the relative size of the *C elegans* germline, it is possible that activity of such cytoplasmic poly(A) polymerases may influence global poly(A) tail length distributions.

Finally, we have created a custom track hub for exploration of this dataset by independent researchers. By making this data easily accessible, we hope to provide *C. elegans* researchers with information related to their genes of interest, providing a resource to identify what isoforms have full-length support in any given developmental stage, and across all stages, as well as the structure of any 3’UTRs that we identify. Given that our dataset provides support for over 7000 isoforms previously lacking full-length support and over 20,000 splice isoforms overall, and given that most isoforms have an associated 3’UTR called, this will be an excellent resource for the *C elegans* research community. Overall, we have demonstrated the utility of nanopore sequencing in providing support for full-length transcripts, annotating putative 3’UTRs, and interrogating poly(A) tail lengths.

## Methods

### *C. elegans* strains, maintenance, and collection

*C. elegans* N2 worms were grown and maintained under standard laboratory conditions on NGM plates seeded with *E. coli* OP50 (Stiernagle 2006). Samples for RNA analysis were synchronized by hypochlorite treatment and overnight hatching in M9 buffer. They were plated as L1d at 25°C and staged by pharyngeal pumping. L2, L3, L4 and young adult (YA) worms were collected approximately two hours post-lethargus. L1 worms were collected four hours after plating. Mature adults were collected approximately ten hours post-L4/YA transition. CB1489 [*him-8(e1489)*IV] adult males were enriched by filtering through 35um mesh.

### RNA extraction

Total RNA isolation was performed using TriReagent (Ambion) following the vendor’s protocol, with the following alterations: three rounds of freeze/thaw lysis were conducted prior to the addition of BCP; RNA was precipitated in isopropanol supplemented with glycogen for one hour at −80°C; RNA was pelleted by centrifugation at 4°C for 30 min at 20,000 x g; the pellet was washed three times in 70% ethanol; the pellet was resuspended in water.

### Library preparation and sequencing

Approximately 20 µg aliquots of total RNA were diluted to a total volume of 100 µl in nuclease free water and poly-A selected using NEXTflex Poly(A) Beads (BIOO Scientific Cat#NOVA-512980). Up to 600 ng of the resulting poly-A RNA was separately aliquoted for library generation. Any excess poly-A selected RNA was stored at −80°C. Biological poly-A RNA and a synthetic control (Lexogen SIRV Set 3, 2.5 ng) were prepared for nanopore direct RNA sequencing generally following the ONT SQK-RNA001 kit protocol, including the optional reverse transcription step recommended by ONT. One difference from the standard ONT protocol was use of Superscript IV (Thermo Fisher) for reverse transcription. RNA sequencing on the GridION platform was performed using ONT R9.4 flow cells and the standard MinKNOW protocol script (NC_48Hr_sequencing_FLO-MIN106_SQK-RNA001).

### Preprocessing and alignments

Reads were basecalled and trimmed of adapter sequences using Poreplex version 0.3.1 (running Albacore version 2.3.1) with the following parameters: -p 24 --trim-adapter --basecall (https://github.com/hyeshik/poreplex). For each of our samples, reads were aligned to the WBcel235 ce11 genome using minimap2 version 2.14-r883 (Li 2018). Genomic alignments were run with the following parameters: -ax splice -k14 -uf --secondary=no -G 25000 -t 24. The resulting.sam files were converted to bam format using samtools view with parameters -b -F 2048 (Li et al. 2009).

### Read filtering

Our first filtering step involved removing reads aligning to the genome with large insertions (>20bp) and large 3’ softclips (>20bp) that could be the result of not properly aligning internal or 3’ exons respectively. This filtering step ensures that novel isoforms identified in downstream scripts are not false positives resulting from poor alignments.

Following this, reads were filtered based on their QC tags from the polyA estimation module of the program nanopolish (Workman et al. 2018). Reads were removed from consideration if they had QC tags “READ_FAILED_LOAD”, “SUFFCLIP”, or “NOREGION”. This was meant to remove reads without a detectable poly(A) tail signal, to prevent inclusion of reads with truncated 3’ ends.

Next, for the purposes of better identifying 3’UTR isoforms in downstream analysis, 3’ soft-clips were realigned using a semi-global aligner with affine gap penalties anchored at the 3’ end of the original alignment. This resulted in more uniform 3’ ends of alignment. The resulting realigned reads were converted to bed12 format using the bedtools bamtobed function (version 2.27.1) (Quinlan 2014; Quinlan and Hall 2010).

To identify full length reads we made use of the Wormbase (release WS265) gff3 gene annotation file (WormBase web site 2018). We converted the Wormbase coding sequence annotations in this file to bed format using a custom python script, resulting in a cds.bed file. We then intersected the cds.bed file with the bed files describing our genomic alignments using the bedtools intersect function, with the flags -s -F 1.0 -u, which enforces that reads span the full length of an annotated CDS (with the correct strandedness) in order to be considered a full-length read. Following this, we collected the read IDs of the resulting full-length reads, and used these IDs to filter genomic alignment files in to full-length and non full-length reads using custom python scripts utilizing pysam (https://github.com/pysam-developers/pysam) version 0.14.1 (Li et al. 2009).

Reads were then filtered to ensure their 5’ ends were within 100 bp of an annotated transcript start site in the Wormbase WS265 gff3 file. This was meant to further reduce the impact of 5’ truncated reads.

To account for errors in splice junction alignments, we used the Wormbase WS265 gff3 annotation to define canonical donor and acceptor splice sites, and assigned each donor and acceptor splice site in our reads to a canonical splice site. Non-canonical donor and acceptor splice sites in our reads that fell within 15 bp of a canonical site were assigned to that site. Reads that contained non-canonical donor and acceptor splice sites that were not within 15 bp of a canonical site were thrown out, and not considered for the purposes of defining splice isoforms or UTRs. In addition, reads were thrown out if splice junctions in that read corresponded to annotated splice junctions from more than one gene. This allowed us to unambiguously assign each spliced read to a gene based on its correspondence to annotated donor and acceptor splice sites. Reads were assigned to splice isoforms in a similar manner (however some of these assignments were ambiguous when two annotated isoforms were comprised of the same sets of splice junctions). For non spliced reads, we assign gene ids based on overlap with single exon genes present in the annotation.

Finally, we separated reads that had exons that span the full length of any intron in the annotation that is not fully spanned by an exon in the annotation. We do this to remove intron retention transcripts from consideration in defining putative isoforms, as we believe these reads to be nuclear RNA that has not fully been processed, which, if included would artificially inflate the number of identified isoforms. Intron retention reads are considered in analysis of poly(A) tail length distributions, as well as in the comparison of poly(A) tail length distributions in fully spliced versus intron retention transcripts.

Reads were excluded from consideration in 3’UTR calling (but not splice isoform calling) if their original minimap2 alignments had 3’ softclips larger than 10 bases long. This exclusion prevented reads with 3’ ends that failed to align well from being considered, and reduced the variation in considered 3’ alignment ends significantly.

### Splice isoform identification

After these stringent filtering steps, we extracted the sequences from the ce11 WBcel235 genome corresponding to each aligned read using getfasta function of the program bedtools with the following flags -s -split -bedOut (Quinlan 2014). We then clustered reads (and their associated sequences) together in to putative isoforms if the reads shared a common set of splice junctions. This resulted in reads clustered by splice isoform. For each of these sets of reads corresponding to splice isoforms, we chose a representative read by selecting the longest read. From this representative read, we extracted information about the isoform including putative coding sequence by identifying the longest open reading frame (with both start and stop codons) present in the read’s associated sequence. This allowed us to define putative start and stop codons.

Splice isoforms were called as novel if they contained a set of splice junctions not previously annotated in the reference. To deal with the possibility of 5’ truncated reads artificially inflating our novel isoform counts, we considered all possible 5’ truncations of previously annotated transcripts in the WormBase WS265 annotation file when defining our reference.

### 3’ UTR calling

To identify putative 3’UTRs, reads were first grouped by their putative stop codons and any splice junctions that occurred downstream of that stop codon. For each read in each of these groups the 3’ most base in their alignment was extracted. These end positions were then used to generate a gaussian kernel density estimate (using the python package seaborn, version 0.9.0 kdeplot function with a specified kernel width of 10). Local maxima in this kernel density estimate were identified, and reported as a putative 3’UTR cleavage site if there were at least 3 read end positions within 10 bp of that local maxima. Reads were assigned to a given 3’UTR if that UTR’s putative cleavage site was the closest UTR cleavage site to the end position of the read, and if the end position of the read and the putative cleavage site were within 10 bp of each other.

### Poly(A) tail length estimation

Poly(A) tail lengths were estimated from raw signal for each read using the polya function of the program nanopolish (version 0.10.2) (Workman et al. 2018). Poly(A) tail length estimates were only considered if the QC tag reported by nanopolish was PASS. Poly(A) tail length estimates were grouped by gene and isoform using the gene and isoform assignments for each read derived from comparison of genomic alignments with the splice junctions in the Wormbase WS265 gff3 reference.

### Calculating coverage for the metagene plot

To generate the metagene plot displayed in Figure 1B, we calculated coverage across every gene (as defined by the ce11 WB245 wormbase .gtf annotation file converted to bed format) using the pybedtools coverage function (Dale et al. 2011; Quinlan and Hall 2010; Lee et al. 2018). We then summed these coverage values together, and normalized the resulting values by dividing each value by the sum of all the coverage values. Genes sizes were scaled such that the size of the gene body and the UTRs were always the same.

### Determining full length support from WormBase annotations

A WormBase splice isoform was said to have full length support if every one of its introns in the WS265 annotation gff3 was annotated to have support from the same EST or the same cDNA (Lee et al. 2018; WormBase web site 2018). This restricted our analysis to only consider isoforms that were annotated as having introns, and excluded single exon genes and genes without introns annotated in the gff3 annotation file (which includes all non-coding RNAs). To account for this, when comparing the number of genes and isoforms we support to the number of genes and isoforms with full length support in WormBase, we only considered splice isoforms from our dataset that corresponded to an isoform from the restricted WormBase isoform set.

### 3’UTR comparisons

We compared our 3’UTRs to the 3’UTRs identified in Jan et al and Mangone et al using a custom script that required putative stop codons match identically, but allowed for a 10 bp tolerance in putative 3’UTR end positions (Jan et al. 2011; Mangone et al. 2010). We identified novel 3’UTRs in a similar manner, but also added consideration of WormBase annotated 3’UTRs.

### Calling PAS sites

We identified PAS sites in a method similar to that used by Mangone et al, in which we searched the 60 nucleotides upstream of the putative cleavage site for putative PAS hexamers (Mangone et al. 2010). Rather than recalculating the frequency of putative PAS hexamers upstream of our putative cleavage sites, we used the PAS hexamers specified in Table S5 of Mangone et al and searched for these hexamers in the order they appear in that Table. Once a putative PAS site was identified, the UTR was assigned that PAS hexamer. If the 3’UTR had none of the hexamers present in the table in it’s upstream sequence, the UTR was said to have no PAS.

### Plotting PAS nucleotide distributions

To plot the nucleotide distribution around a given type of PAS site, we first sorted sequences by their PAS type. For canonical and alternative PAS sites, nucleotide distributions were anchored such that the PAS site began at −19 nucleotides. The percentage of use of each base at each position in a window around the PAS site was then calculated. For UTRs with no PAS identified, the nucleotide distribution was calculated such that the putative cleavage site was at position 0.

## Software availability

Code required to replicate the analyses performed in this paper is available on GitHub at https://github.com/NatPRoach/c_elegans_dRNAseq_analysis.

## Data Access

Both raw fast5 and basecalled fastq have been deposited at the European Nucleotide Archive (ENA) and can be found under accession number PRJEB31791.

## Acknowledgements

This work was supported by a grant from the NIH to JKK. (NIH R01GM118875) and by a Johns Hopkins Discovery Award to WT, JT, and JKK. NPR was partly supported by a training grant awarded to the Johns Hopkins Cell, Molecular, Developmental Biology and Biophysics program (NIH T32GM007231). We thank Mindy Clark for the *C. elegans* life cycle diagram in Figure 1A. We thank Mallory Freeberg for initial computational analyses comparing nanopore based cDNA and dRNA sequencing that led us to utilize dRNAseq in this study.

## Author contributions statement

AA collected the developmental samples for *C. elegans* and isolated the RNA to be sequenced and NS performed the sequencing. NR performed all of the sequencing analysis in consultation with J.T.. All authors reviewed the manuscript.

## Supplemental Materials

**Supplemental Figure 1.**
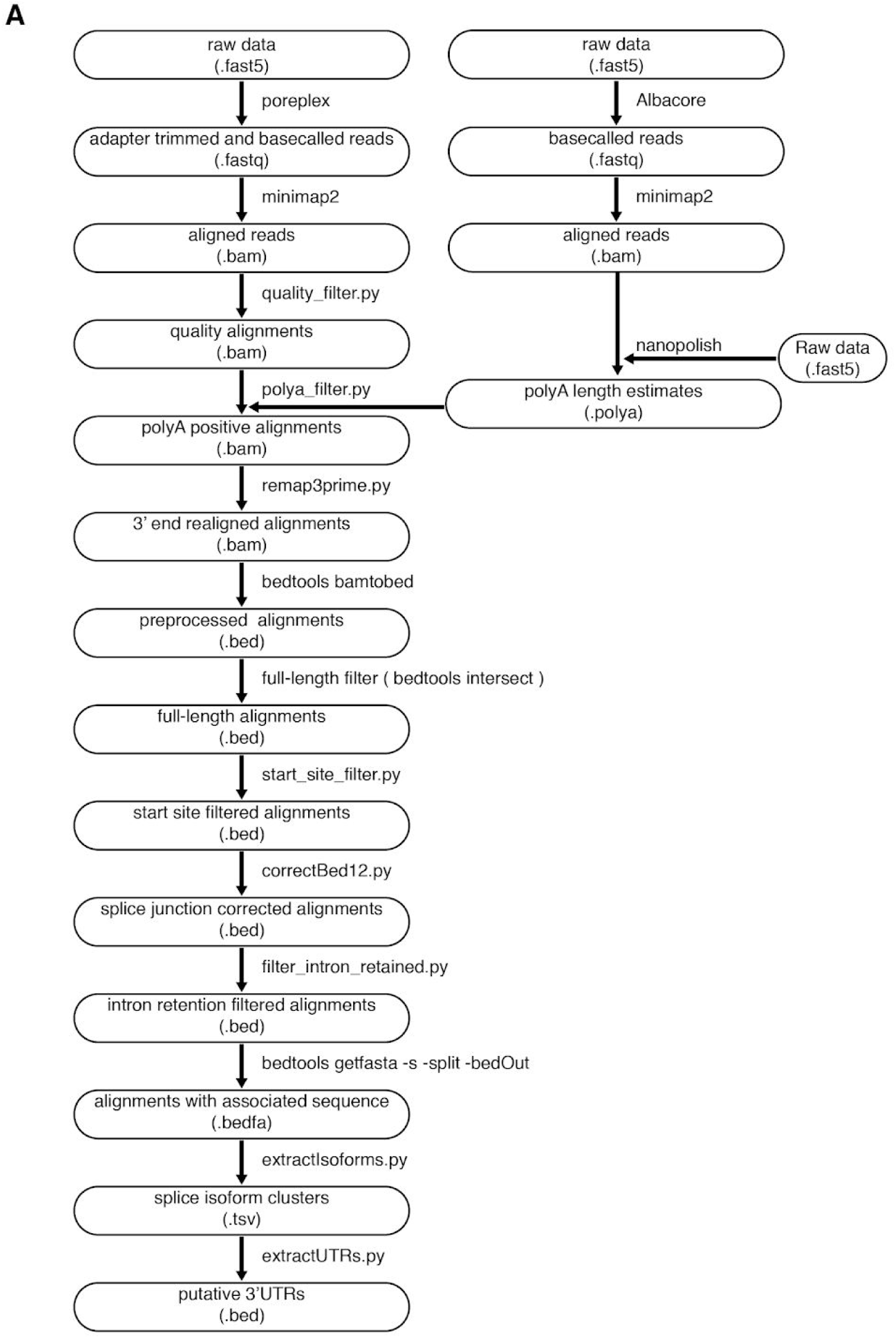
Flowchart of computational analysis.

**Supplemental Figure 2.**
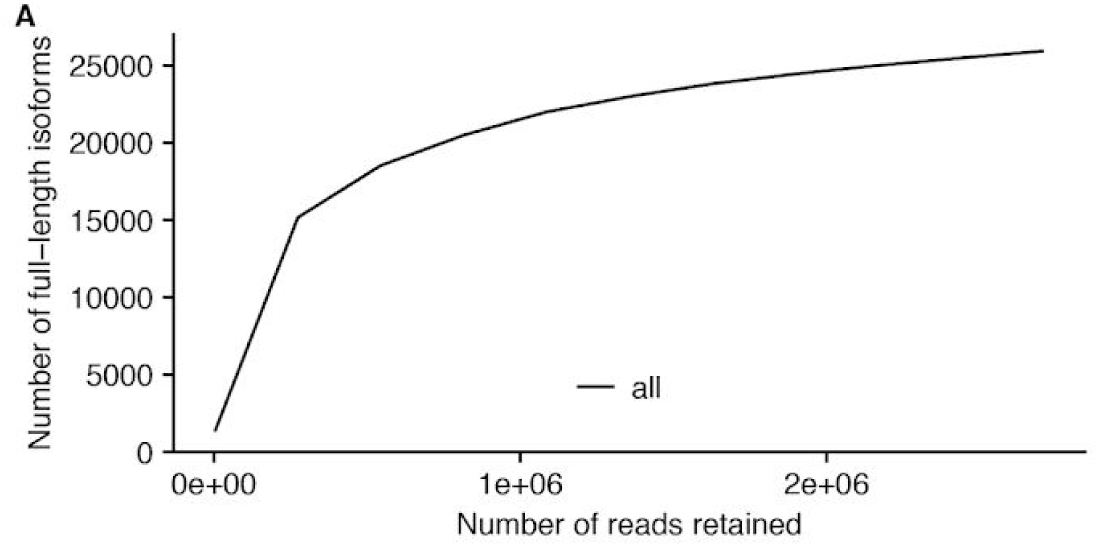
Saturation plot of full length isoforms identified in all stages pooled (as in Figure 1G).

**Supplemental Figure 3.**
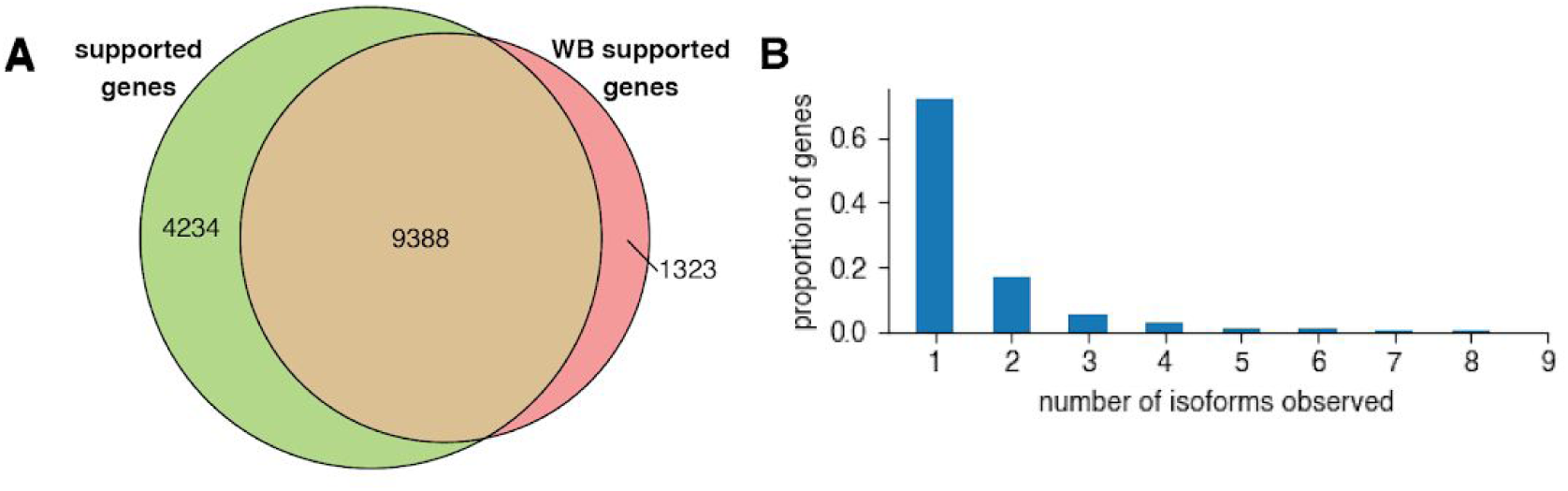
**(A)** Venn diagram showing overlap between genes identified with full-length support in our dataset, and genes with full length support in the WormBase (WB) dataset. **(B)** Density plot showing distribution of number of isoforms observed per gene in the Wormbase WS265 gff3 annotation. (WormBase web site 2018; Lee et al. 2018).

**Supplemental Figure 4.**
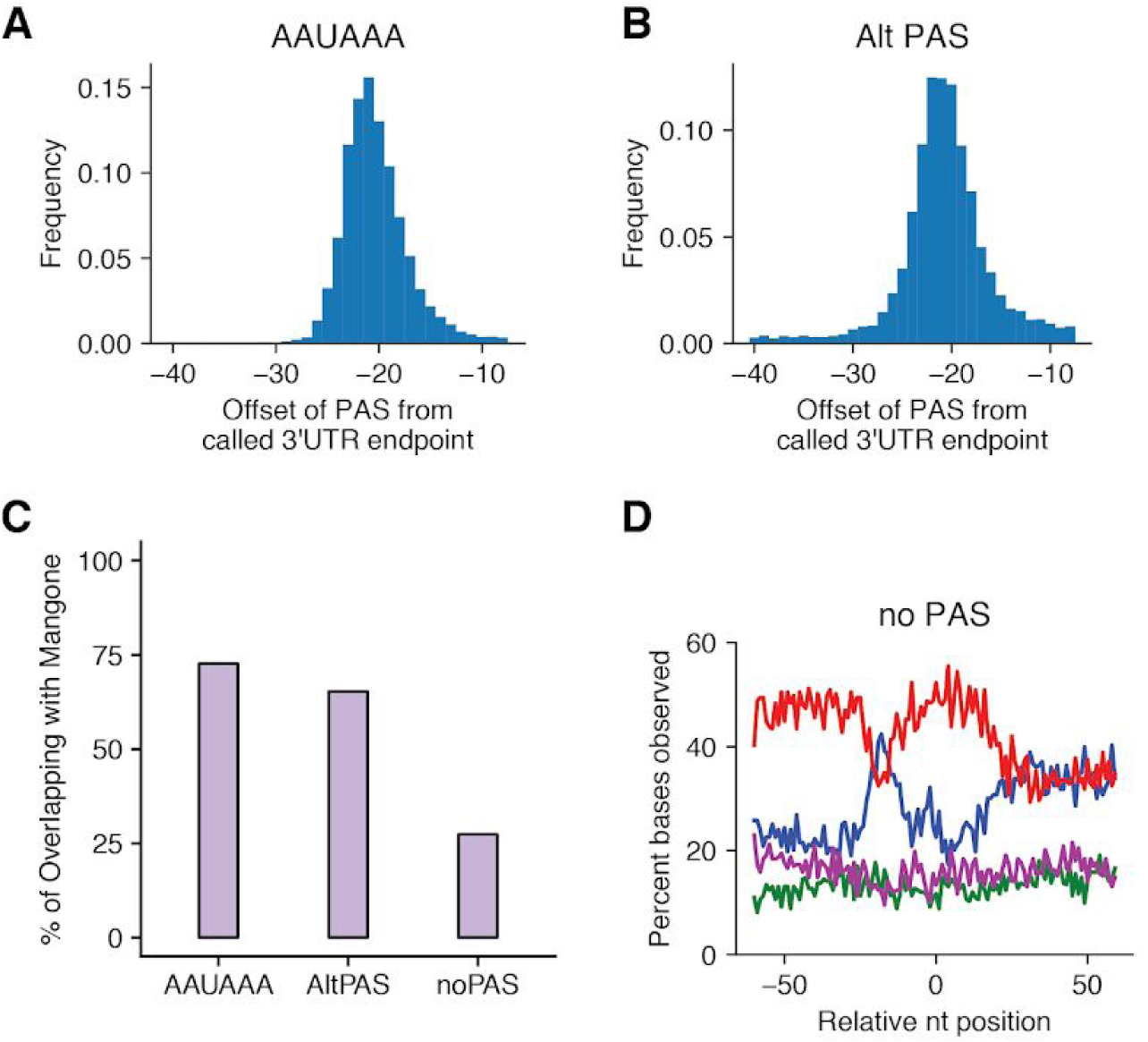
Evidence supporting the validity of our identified 3’UTRs. Offsets of identified PAS sites from the putative cleavage site for canonical **(A)** and non-canonical **(B)** PAS sites. **(C)** Percent of UTRs with specified PAS site type that overlap with a Mangone et al 3’UTR. **(D)** Nucleotide distribution in a window around putative cleavage sites for 3’UTRs that overlap with a Mangone 3’UTR and do not have a PAS site identified. This distribution is different than the published distribution of no PAS Mangone 3’UTRs in general (Mangone et al. 2010).

**Supplemental Figure 5.**
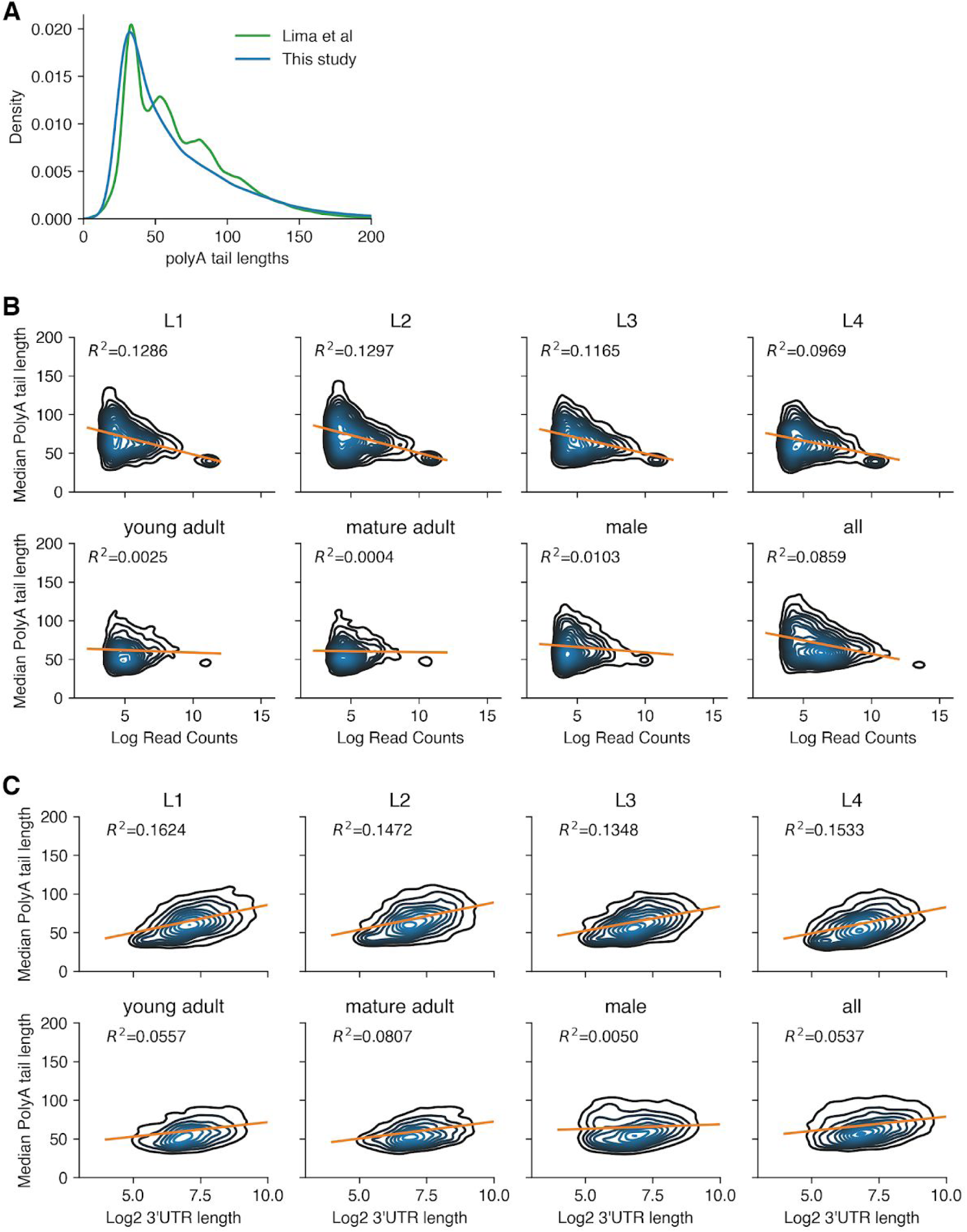
Comparison of poly(A) tail length distributions between reads from our L4 stage dataset and Lima et al(Lima et al. 2017). **(A)**. Density plots including linear regressions (orange line) of median poly(A) tail length versus expression level **(B)** or 3’UTR length **(C)**, separated by stage.

## References

Andreassi C, Riccio A. 2009. To localize or not to localize: mRNA fate is in 3’ UTR ends. Trends Cell Biol 19: 465–474.

Bayega A, Oikonomopoulos S, Zorbas E, Wang YC, Gregoriou M-E, Tsoumani KT, Mathiopoulos KD, Ragoussis J. 2018. Transcriptome landscape of the developing olive fruit fly embryo delineated by Oxford Nanopore long-read RNA-Seq. bioRxiv 478172. https://www.biorxiv.org/content/10.1101/478172v1.abstract (Accessed March 11, 2019).

Blazie SM, Geissel HC, Wilky H, Joshi R, Newbern J, Mangone M. 2017. Alternative Polyadenylation Directs Tissue-Specific miRNA Targeting in Caenorhabditis elegans Somatic Tissues. Genetics 206: 757–774.

Broverman SA, Meneely PM. 1994. Meiotic mutants that cause a polar decrease in recombination on the X chromosome in Caenorhabditis elegans. Genetics 136: 119–127.

Byrne A, Beaudin AE, Olsen HE, Jain M, Cole C, Palmer T, DuBois RM, Forsberg EC, Akeson M, Vollmers C. 2017. Nanopore long-read RNAseq reveals widespread transcriptional variation among the surface receptors of individual B cells. Nat Commun 8: 16027.

Cai Y, Yu X, Hu S, Yu J. 2009. A Brief Review on the Mechanisms of miRNA Regulation. Genomics Proteomics Bioinformatics 7: 147–154.

Chang H, Lim J, Ha M, Kim VN. 2014. TAIL-seq: genome-wide determination of poly(A) tail length and 3’ end modifications. Mol Cell 53: 1044–1052.

Corsi AK, Wightman B, Chalfie M. 2015. A Transparent Window into Biology: A Primer on Caenorhabditis elegans. Genetics 200: 387–407.

Dale RK, Pedersen BS, Quinlan AR. 2011. Pybedtools: a flexible Python library for manipulating genomic datasets and annotations. Bioinformatics 27: 3423–3424.

Diag A, Schilling M, Klironomos F, Ayoub S, Rajewsky N. 2018. Spatiotemporal m(i)RNA Architecture and 3’ UTR Regulation in the C. elegans Germline. Developmental Cell 47: 785–800.e8. http://dx.doi.org/10.1016/j.devcel.2018.10.005.

Garalde DR, Snell EA, Jachimowicz D, Sipos B, Lloyd JH, Bruce M, Pantic N, Admassu T, James P, Warland A, et al. 2018. Highly parallel direct RNA sequencing on an array of nanopores. Nat Methods 15: 201–206.

Gerstein MB, Lu ZJ, Van Nostrand EL, Cheng C, Arshinoff BI, Liu T, Yip KY, Robilotto R, Rechtsteiner A, Ikegami K, et al. 2010. Integrative analysis of the Caenorhabditis elegans genome by the modENCODE project. Science 330: 1775–1787.

Hillier LW, Coulson A, Murray JI, Bao Z, Sulston JE, Waterston RH. 2005. Genomics in C. elegans: so many genes, such a little worm. Genome Res 15: 1651–1660.

Hillier LW, Reinke V, Green P, Hirst M, Marra MA, Waterston RH. 2009. Massively parallel sequencing of the polyadenylated transcriptome of C. elegans. Genome Res 19: 657–666.

Hodgkin J, Horvitz HR, Brenner S. 1979. Nondisjunction Mutants of the Nematode CAENORHABDITIS ELEGANS. Genetics 91: 67–94.

Jan CH, Friedman RC, Ruby JG, Bartel DP. 2011. Formation, regulation and evolution of Caenorhabditis elegans 3’UTRs. Nature 469: 97–101.

Jenjaroenpun P, Wongsurawat T, Pereira R, Patumcharoenpol P, Ussery DW, Nielsen J, Nookaew I. 2018. Complete genomic and transcriptional landscape analysis using third-generation sequencing: a case study of Saccharomyces cerevisiae CEN.PK113-7D. Nucleic Acids Res 46: e38.

Kadobianskyi M, Schulze L, Schuelke M, Judkewitz B. 2019. Hybrid genome assembly and annotation of Danionella translucida, a transparent fish with the smallest known vertebrate brain. bioRxiv 539692. https://www.biorxiv.org/content/10.1101/539692v1.abstract (Accessed April 2, 2019).

Kuersten S, Goodwin EB. 2003. The power of the 3’ UTR: translational control and development. Nat Rev Genet 4: 626–637.

Lamesch P, Milstein S, Hao T, Rosenberg J, Li N, Sequerra R, Bosak S, Doucette-Stamm L, Vandenhaute J, Hill DE, et al. 2004. C. elegans ORFeome version 3.1: increasing the coverage of ORFeome resources with improved gene predictions. Genome Res 14: 2064–2069.

Lee RC, Feinbaum RL, Ambros V. 1993. The C. elegans heterochronic gene lin-4 encodes small RNAs with antisense complementarity to lin-14. Cell 75: 843–854.

Lee RYN, Howe KL, Harris TW, Arnaboldi V, Cain S, Chan J, Chen WJ, Davis P, Gao S, Grove C, et al. 2018. WormBase 2017: molting into a new stage. Nucleic Acids Res 46: D869–D874.

Legnini I, Alles J, Karaiskos N, Ayoub S, Rajewsky N. 2019. Full-length mRNA sequencing reveals principles of poly (A) tail length control. bioRxiv. https://www.biorxiv.org/content/10.1101/547034v1.abstract.

Li H. 2018. Minimap2: pairwise alignment for nucleotide sequences. Bioinformatics 34: 3094–3100.

Li H, Handsaker B, Wysoker A, Fennell T, Ruan J, Homer N, Marth G, Abecasis G, Durbin R, 1000 Genome Project Data Processing Subgroup. 2009. The Sequence Alignment/Map format and SAMtools. Bioinformatics 25: 2078–2079.

Lima SA, Chipman LB, Nicholson AL, Chen Y-H, Yee BA, Yeo GW, Coller J, Pasquinelli AE. 2017. Short poly(A) tails are a conserved feature of highly expressed genes. Nat Struct Mol Biol 24: 1057–1063.

Lim J, Lee M, Son A, Chang H, Kim VN. 2016. mTAIL-seq reveals dynamic poly(A) tail regulation in oocyte-to-embryo development. Genes Dev 30: 1671–1682.

Mangone M, Manoharan AP, Thierry-Mieg D, Thierry-Mieg J, Han T, Mackowiak SD, Mis E, Zegar C, Gutwein MR, Khivansara V, et al. 2010. The Landscape of C. elegans 3’UTRs. Science 329: 432–435.

Mayr C, Bartel DP. 2009. Widespread shortening of 3’ UTRs by alternative cleavage and polyadenylation activates oncogenes in cancer cells. Cell 138: 673–684.

Millonigg S, Minasaki R, Nousch M, Eckmann CR. 2014. GLD-4-Mediated Translational Activation Regulates the Size of the Proliferative Germ Cell Pool in the Adult C. elegans Germ Line. PLoS Genetics 10: e1004647. http://dx.doi.org/10.1371/journal.pgen.1004647.

Nousch M, Minasaki R, Eckmann CR. 2017. Polyadenylation is the key aspect of GLD-2 function in C. elegans. RNA 23: 1180–1187.

Packer JS, Zhu Q, Huynh C, Sivaramakrishnan P. 2019. A lineage-resolved molecular atlas of C. elegans embryogenesis at single cell resolution. bioRxiv. https://www.biorxiv.org/content/10.1101/565549v2.abstract.

Pertea M, Pertea GM, Antonescu CM, Chang T-C, Mendell JT, Salzberg SL. 2015. StringTie enables improved reconstruction of a transcriptome from RNA-seq reads. Nat Biotechnol 33: 290–295.

Phillips CM, Wong C, Bhalla N, Carlton PM, Weiser P, Meneely PM, Dernburg AF. 2005. HIM-8 binds to the X chromosome pairing center and mediates chromosome-specific meiotic synapsis. Cell 123: 1051–1063.

Quinlan AR. 2014. BEDTools: the Swiss-army tool for genome feature analysis. Curr Protoc Bioinformatics 47: 11–12.

Quinlan AR, Hall IM. 2010. BEDTools: a flexible suite of utilities for comparing genomic features. Bioinformatics 26: 841–842.

Reboul J, Vaglio P, Tzellas N, Thierry-Mieg N, Moore T, Jackson C, Shin-i T, Kohara Y, Thierry-Mieg D, Thierry-Mieg J, et al. 2001. Open-reading-frame sequence tags (OSTs) support the existence of at least 17,300 genes in C. elegans. Nat Genet 27: 332–336.

Saito TL, Hashimoto S-I, Gu SG, Morton JJ, Stadler M, Blumenthal T, Fire A, Morishita S. 2013. The transcription start site landscape of C. elegans. Genome Res 23: 1348–1361.

Schmid M, Küchler B, Eckmann CR. 2009. Two conserved regulatory cytoplasmic poly(A) polymerases, GLD-4 and GLD-2, regulate meiotic progression in C. elegans. Genes Dev 23: 824–836.

Sessegolo C, Cruaud C, Da Silva C, Dubarry M, Derrien T, Lacroix V, Aury J-M. 2019. Transcriptome profiling of mouse samples using nanopore sequencing of cDNA and RNA molecules. bioRxiv 575142. https://www.biorxiv.org/content/10.1101/575142v1.abstract (Accessed April 2, 2019).

Soneson C, Yao Y, Bratus-Neuenschwander A, Patrignani A, Robinson MD, Hussain S. 2019. A comprehensive examination of Nanopore native RNA sequencing for characterization of complex transcriptomes. http://dx.doi.org/10.1101/574525.

Spieth J, Genome Sequencing Center, Washington University School of Medicine, Louis S, Usa MO 63108. 2014. Overview of gene structure in C. elegans. WormBook 1–18. http://dx.doi.org/10.1895/wormbook.1.65.2.

Stiernagle T. 2006. Maintenance of C. elegans. WormBook. The C. elegans research community. WormBook.

Subtelny AO, Eichhorn SW, Chen GR, Sive H, Bartel DP. 2014. Poly(A)-tail profiling reveals an embryonic switch in translational control. Nature 508: 66–71. http://dx.doi.org/10.1038/nature13007.

Suh N, Jedamzik B, Eckmann CR, Wickens M, Kimble J. 2006. The GLD-2 poly (A) polymerase activates gld-1 mRNA in the Caenorhabditis elegans germ line. Proceedings of the National Academy of Sciences 103: 15108–15112.

Sulston JE, Schierenberg E, White JG, Thomson JN. 1983. The embryonic cell lineage of the nematode Caenorhabditis elegans. Dev Biol 100: 64–119.

Szostak E, Gebauer F. 2013. Translational control by 3’-UTR-binding proteins. Brief Funct Genomics 12: 58–65.

Tang AD, Soulette CM, van Baren MJ, Hart K. 2018. Full-length transcript characterization of SF3B1 mutation in chronic lymphocytic leukemia reveals downregulation of retained introns. bioRxiv. https://www.biorxiv.org/content/10.1101/410183v1.abstract.

The C. elegans Sequencing Consortium. 1998. Genome Sequence of the Nematode C. elegans: A Platform for Investigating Biology. Science 282: 2012–2018. http://dx.doi.org/10.1126/science.282.5396.2012.

Tourasse NJ, Millet JRM, Dupuy D. 2017. Quantitative RNA-seq meta-analysis of alternative exon usage in C. elegans. Genome Res 27: 2120–2128.

Trapnell C, Roberts A, Goff L, Pertea G, Kim D, Kelley DR, Pimentel H, Salzberg SL, Rinn JL, Pachter L. 2012. Differential gene and transcript expression analysis of RNA-seq experiments with TopHat and Cufflinks. Nat Protoc 7: 562–578.

Volden R, Palmer T, Byrne A, Cole C, Schmitz RJ, Green RE, Vollmers C. 2018. Improving nanopore read accuracy with the R2C2 method enables the sequencing of highly multiplexed full-length single-cell cDNA. Proceedings of the National Academy of Sciences 115: 9726–9731. http://dx.doi.org/10.1073/pnas.1806447115.

Walhout AJM, Temple GF, Brasch MA, Hartley JL, Lorson MA, van den Heuvel S, Vidal M. 2000. [34] GATEWAY recombinational cloning: Application to the cloning of large numbers of open reading frames or ORFeomes. In Methods in Enzymology (eds. J. Thorner, S.D. Emr, and J.N. Abelson), Vol. 328 of, pp. 575–IN7, Academic Press.

Wightman B, Bürglin TR, Gatto J, Arasu P, Ruvkun G. 1991. Negative regulatory sequences in the lin-14 3’-untranslated region are necessary to generate a temporal switch during Caenorhabditis elegans development. Genes Dev 5: 1813–1824.

Wightman B, Ha I, Ruvkun G. 1993. Posttranscriptional regulation of the heterochronic gene lin-14 by lin-4 mediates temporal pattern formation in C. elegans. Cell 75: 855–862.

Williams GW, Davis PA, Rogers AS, Bieri T, Ozersky P, Spieth J. 2011. Methods and strategies for gene structure curation in WormBase. Database 2011: baq039.

Wilson RK. 1999. How the worm was won. The C. elegans genome sequencing project. Trends Genet 15: 51–58.

Workman RE, Tang A, Tang PS, Jain M, Tyson JR. 2018. Nanopore native RNA sequencing of a human poly (A) transcriptome. BioRxiv. https://www.biorxiv.org/content/10.1101/459529v1.abstract.

WormBase web site. 2018. http://wormbase.org, release WS265.

Zerbino DR, Achuthan P, Akanni W, Amode MR, Barrell D, Bhai J, Billis K, Cummins C, Gall A, Girón CG, et al. 2018. Ensembl 2018. Nucleic Acids Res 46: D754–D761.

Zhao H-Q, Zhang P, Gao H, He X, Dou Y, Huang AY, Liu X-M, Ye AY, Dong M-Q, Wei L. 2015. Profiling the RNA editomes of wild-type C. elegans and ADAR mutants. Genome Res 25: 66–75.

